# ThermoLink: Bridging Disulfide Bond and Enzyme Thermostability through Database Construction and Machine Learning Prediction

**DOI:** 10.1101/2023.11.30.569109

**Authors:** Ran Xu, Yilin Ye, Minghui Xin, Zechen Wang, Sheng Wang, Weifeng Li, Yanjie Wei, Liangzhen Zheng, Jingjing Guo

## Abstract

Disulfide bonds (SS bonds), covalently formed by sulfur atoms in cysteine residues, play a crucial role in protein folding and structure stability. Due to their significance, artificial disulfide bonds are often introduced to enhance the thermostability of proteins. Although an increasing number of tools can assist with this task, significant amounts of time and resources were often wasted due to inadequate consideration. To enhance the accuracy and efficiency of designing disulfide bonds for protein thermostability improvement, we first collected disulfide-bond data with protein thermostability data from extensive literature sources. Then, various sequence- and structure-based features were extracted, and machine learning models were constructed to predict whether a disulfide bond could improve protein thermostability. Among all models, the neighborhood context model using the Adaboost-DT algorithm performs the best, and the AUC-ROC score and accuracy are 0.773 and 0.714, respectively. Alongside this, we also found that AlphaFold2 exhibits a high superiority in predicting disulfide bonds, and the coevolutionary relationship between residue pairs, to some extent, could also guide artificial disulfide-bond design. The SS-bond data has been integrated into an online server, named ThermoLink, available at guolab.mpu.edu.mo/thermoLink.

Key Messages

- We manually curated a database of disulfide bonds and their impacts on protein thermostability from the literature.
- AlphaFold2 exhibits a high superiority in predicting disulfide bonds, and to some extent, the conservation and co-evolutionary information of the residue pairs involved in disulfide bonds could also guide artificial disulfide-bond design.
- Machine learning models were developed to predict potential disulfide bonds with the aim of improving protein thermostability, offering an extended design strategy beginning with existing SS-bond prediction tools to optimize variant selection.
- This work provides valuable data and theoretical support for a more precise design of disulfide bonds for improving protein thermostability.

## Introduction

Thermostability is an essential requirement for industrial enzymes, as it allows efficient reactions at high temperatures and prolongs the enzyme half-life[1, 2]. Although ongoing improvement opportunities exist for enzyme activity and stability, natural evolution does not guarantee their optimal states in terms of both thermostability and activity. This is because enzymes evolve in response to changing environmental conditions and selective pressures, which may prioritize certain aspects of enzyme function over others[3]. As a result, enzymes may possess inherent trade-offs or suboptimal characteristics that limit their overall performance. Through the evolution of the direction[4] or engineering[5, 6], it is possible to improve the performance of enzymes by enhancing their catalytic efficiency, substrate specificity, resistance to denaturation, and tolerance to harsh conditions (such as a high concentration of organic molecules) for industrial applications[7, 8]. This optimization process involves modifying key amino acids by introducing beneficial mutations, short peptide tags, or sometimes large sequence substitutions to enhance enzyme properties[9, 10, 11]. By iteratively exploring and refining enzyme variants, researchers can unlock the untapped potential within natural enzymes and harness their catalytic power for various industrial, biomedical, and biotechnological applications.

To develop more stable enzymes for industrial applications, various strategies have been proposed and used, such as reverting to consensus mutations, mutating to thermophilic homologs, rigidifying flexible regions, introducing salt or disulfide bridges, and incorporating proline mutations[12, 13, 14, 15, 16]. In particular, the introduction of disulfide bonds (SS bonds) is widely regarded as a viable and effective strategy to improve enzyme thermostability in protein engineering[17]. These covalent bonds, formed between pairs of proximal cysteines, act as structural scaffolds by establishing intra- or inter-molecular connections, reinforce protein structure, and therefore play crucial roles in protein folding, stability, and function[18, 19, 20, 21]. However, whether all of the designed SS bonds would contribute to favorable thermostability improvements has not been strictly quantified.

Now, with the development of structural biology and the emergence of AI-driven tools like AlphaFold2 (AF2)[22], rational protein design is accurately and efficiently facilitated with rich structural information. At the same time, some SS-bond design tools have been developed, including DBD2[23], MODIP[24], Yosshi[25], and so on. However, with the aid of these tools, undesired designs still lead to negative results, including lower thermostability, lower activity, or even loss of function, and so on. Therefore, there is a pressing need to conduct further investigations into the stability increase or functional gain of enzymes when introducing new SS bridges.

Recently, machine learning (ML) has been used to accelerate the evolution of proteins[26, 27]. By training on a large dataset of enzyme sequences and their corresponding characteristics, machine learning algorithms can learn the relationship between them, allowing accurate predictions of the characteristics of new enzyme variants. Different training data can lead to the exploration of different regions of sequence space in ML-guided directed evolution, potentially resulting in varying levels of functional improvement[28]. Through machine learning, it is possible to design new enzyme variants with improved thermostability and functionality by exploring different sequence spaces and SS bond patterns. These machine learning-driven engineering approaches could provide an efficient pathway for developing thermally stable enzymes to address challenges in industrial and biotechnological applications.

Increasing instances of engineered SS bonds have been documented, including successful implementations and failures, as well as undetectable ones[29, 30, 31, 32]. However, the sparse data in the literature can hinder the ability to draw further analyses and robust conclusions, identify patterns, or make accurate predictions. To address the gaps in this rapidly evolving field, we have curated a database named ThermoLink based on a search of a large amount of literature to investigate the specific impact of different types of SS bridges on enzyme thermostability. The sequence- and structure-based features were extracted for in-depth analysis. The exploration of SS bonds and their characteristic in enhancing thermostability has uncovered several intriguing and significant findings. Based on various extracted sequence- and structure-based features, machine learning models were built to predict the thermostability effect of the introduction of an SS bridge. Overall, we have identified patterns and regularities that provide data and theoretical support for a more precise design of SS bonds in the future.

## MATERIALS AND METHODS

### Data collection and processing

#### Data collection

We gathered pertinent data by conducting a thorough literature review and searching reputable online databases such as PubMed, NCBI, UniProt, and other academic sources. Our search criteria included specific keywords and filters to obtain datasets directly relevant to our research question. The collected data was meticulously cleaned and preprocessed to eliminate outliers, address missing values, and ensure data consistency and usability. Details are shown on the left of Fig. 1.

**Figure 1.**
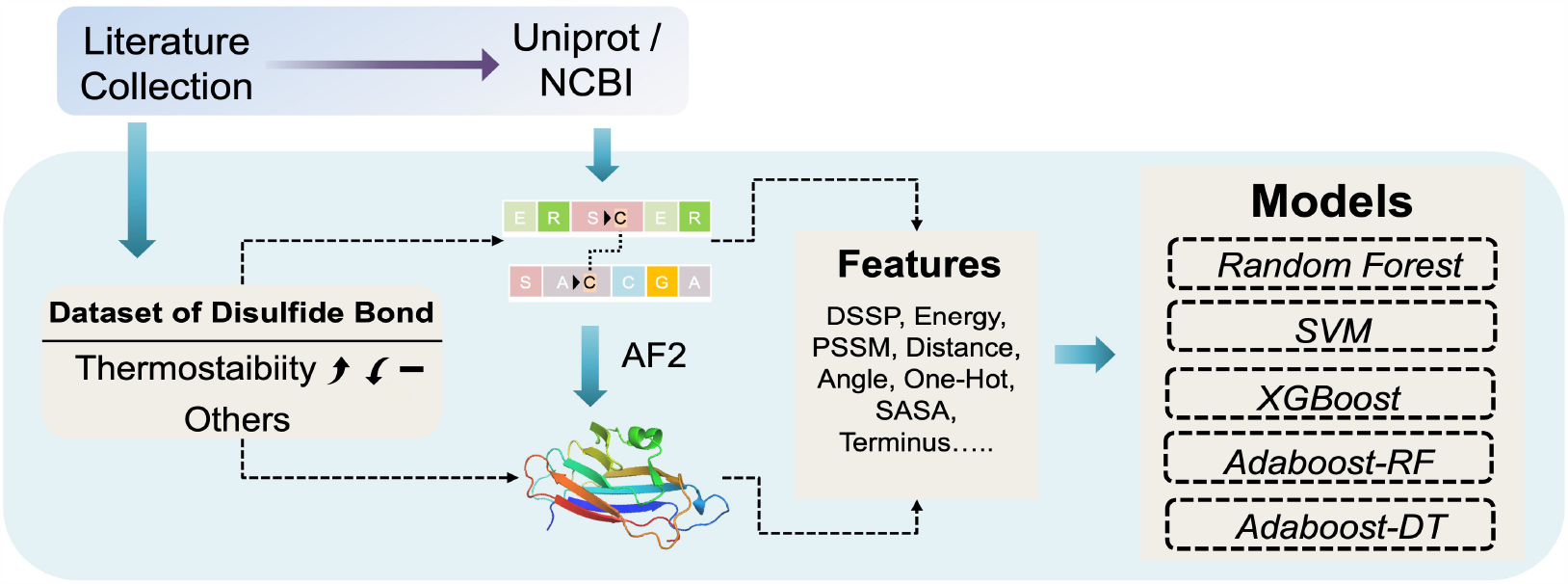
The workflow for this work. Experimental data on the SS bonds that influence enzyme thermostability and function were collected from the literature. The sequence information was derived from the National Center for Biotechnology Information (NCBI) and UniProt databases, and the structures of wild-type and cysteine mutants were predicted by the AI-powered tool, Alphafold2 (AF2). Both sequence- and structure-based features were considered for the construction of the prediction model using multiple machine learning methods, such as Support Vector Machine (SVM), XGBoost, Random Forest (RF), and Adaptive Boosting Decision Tree (Adaboost-DT), Adaptive Boosting Random Forest (Adaboost-RF).

#### Structure prediction using AF2

AF2[22, 33] can provide high-quality predicted 3D protein structures containing more enriched information than 1D amino acid sequences. The sequences collected before and after disulfide-bond modification were adopted for AF2-based structure prediction. AF2 relies on multiple sequence alignment to extract the co-evolution information and structure templates from the Protein Data Bank. Among the predicted five structure models, only the best one (with the highest plDDT score) was used for further analysis. In this study, we adopted the “faster” version of AF2 (fastAF2)[34, 35] for structure prediction, where speed is faster and precision is guaranteed. This fastAF2 has been shown to be helpful in predicting the structure of the protein-ligand complex[36] in the 15th Critical Assessment of Structure Prediction (CASP15).

#### Natural SS-bond structure data acquisition

In this section, PISCES (Protein Sequence Screening Server) was applied to obtain protein chains with less than 50% sequence similarity to each other in the cullpdb database (update as of September 23, 2023)[37, 38]. Then, we extracted the residue numbers of cysteines associated with SS bonds by searching for ”SSBOND” in the PDB files and matching them with the corresponding chain IDs. Finally, 2777 protein chains with intra-chain SS bonds were identified and downloaded.

### Feature extraction

#### Sequence-based feature extraction

Firstly, one-hot encoding[39] is used for protein residue type representations for SS-Bond engineering sites. In addition to focusing on the mutation site itself, the surrounding *N* (where *N* = 5) residues are also taken into account to capture the surrounding environment of the mutation site.

The Position Specific Scoring Matrix (PSSM) has emerged as a piece of valuable information in protein bioinformatics to gain insights into evolutionary conservation and make predictions related to protein folding prediction, RNA binding sites, and protein functional analysis[40, 41, 42]. In this work, PSSM was calculated to evaluate the sequence conservation of the residue pairs forming SS linkages[43]. To collect local information for the specified sites, we perform feature extraction directly on the basis of the PSSM profiles of the designed sites. A higher score indicates a higher likelihood of finding an amino acid at that position, while a lower value indicates a lower likelihood.

#### Structure-based feature extraction

In this work, we calculated the mutation-induced Gibbs free energy change of each residue pair using the FoldX suite[44]. The free energy change of unfolding (Δ*G*) of a target protein and the energy difference between wild type and mutant were calculated using the following equations (1-2)[44]:

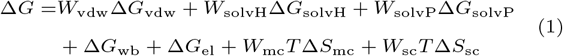

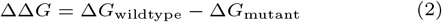

Then, SASA (Solvent Accessible Surface Area)[45], which measures the surface area of atoms accessible to the solvent, has been calculated for residues involved in disulfide bonds using mdtraj[46]. This measurement provides valuable information on the exposure of the residues to the surrounding environment.

Next, the angle and distance between the selected atoms of residues involved in SS bonds have been calculated. These geometric parameters provide information on the spatial arrangement and conformation of the SS bonds. Non-covalent bonds, such as hydrogen bonds and salt bridges, play crucial roles in stabilizing protein structures. Hydrogen bonds and salt bridges formed between residues adjacent to the SS bonds have been identified and quantified to consider polar interactions around the SS bond. The enzyme structures were read using the Biopython module[47], and then the hydrogen bonds and salt bridges formed between the mutation site and the surrounding amino acids were calculated. Hydrogen bonds are defined as distances between hydrogen atoms and oxygen and nitrogen atoms less than the threshold of 3.5 Å, and salt bridges involve interactions between charged residues with a distance cut-off of 4 Å[48]. These interactions may contribute to the overall or partial stability of the protein structure.

Furthermore, the secondary structure information (e.g., *α* helix, *β* strand) of the SS-bond sites and their surrounding residues was calculated using DSSP (dynamic secondary structure of proteins)[49], providing insights into the local structural context of the SS bonds. Eight categories of secondary structures (3_10_–helix (G), *α*–helix (H), *π*–helix (I), *β*–bridge (B), *β*–strand (E), high-curvature loop (S), *β*–turn (T) and coil (C)) obtained from the DSSP calculations were grouped into three main categories: helix (G, H, I); *β*–sheet (B, E); others, (S, T, C and undetected secondary structure).

### Development and validation of machine learning models

#### Training and testing datasets

The collected SS bonds were classified into two groups. Those introducing favorable thermostability were labeled as ”positive”, while others were labeled as ”negative”. The labeled dataset is then used to train a model to predict the effects of SS bonds on protein thermostability. To increase the robustness of the model, the enzymes have been clustered according to a sequence similarity threshold of 0.6 using CD-HIT (version 4.8.1)[50]. The resulting clusters were then randomly divided into training and testing sets for the train-test split (ratio = 9: 1), ensuring that samples from the same cluster were present in the same dataset, either in the training set or the testing set.

#### The construction of machine learning models

To develop a classification model that can differentiate the impact of artificial SS bonds on thermal stability, we utilized the positive and negative sets mentioned in the preceding section. Five supervised algorithms were implemented and trained on those labeled data[51]. These methods include Random forest (RF)[52], XGBoost[53, 54], Adaboost-RF[55], Adaboost-DT[56], and Support vector machine (SVM)[57]. Random Forest, Adaboost-RF, XGBoost, and Adaboost-DT are all ensemble algorithms. These algorithms combine multiple weak learners, such as decision trees, to build a strong learner that improves overall predictive performance. In SVM, a linear kernel is employed to transform the data into a higher-dimensional feature space, enabling the identification of an optimal hyperplane for effective data separation. These machine-learning algorithms have been implemented and integrated into the Python scikit-learn package[58]. The optimal parameters used in our models were obtained via a grid search strategy.

#### Performance evaluation

To verify the model quality, the following parameters were applied:

Accuracy:

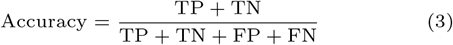

Recall:

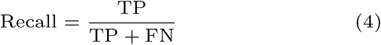

Precision:

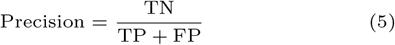

F1 score:

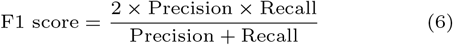

Specificity:

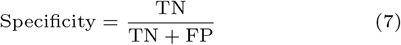

AUC:

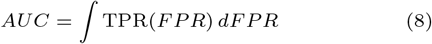

where TP, TN, FP, and FN represent true positives, true negatives, false positives, and false negatives, respectively. Moreover, the area under the receiver operating characteristic (ROC) and precision-recall (PR) curves denoted as AUC were computed, with elevated AUC values signifying enhanced model performances.

## RESULTS

### SS-bond database for enzyme thermostability engineering

In this work, we collected data on SS linkages, both artificial and native, along with corresponding enzyme thermostability information. For each item, we compiled the sequence ID, source, and category of enzymes, as well as the designed SS bond sites and the consequent impact on enzyme thermostability. Within our study, we classify “enhanced thermostability” as positive data (55%), while considering other situations as negative data (Fig. 2A–B). It is worth noting that given the subjectivity of the investigators, there may be many negative samples that were not recorded. Most of the enzymes are derived from bacteria (58%) and fungi (31%), and others include mammals, amphibians, or synthetic constructs (Fig. 2C). According to the IUBMB classification system[59], enzymes can be grouped into six categories: oxidoreductases, transferases, hydrolases, lyases, isomerases, and ligases, which are involved in oxidation-reduction, group transfer, hydrolysis, condensation, isomerization, and ligation reactions, respectively[60, 61]. In our collected data, the proportions of hydrolases and oxidoreductases are quite high, with hydrolases accounting for 60% and oxidoreductases accounting for 32% (Fig. 2D). Hydrolases, as important biological components, play a crucial role in industrial production, including lipases, esterases, amylases, proteases, etc.[62] SS bonds are often introduced into lipases to increase their thermostability, commonly by rigidifying flexible lid domains[63].

**Figure 2.**
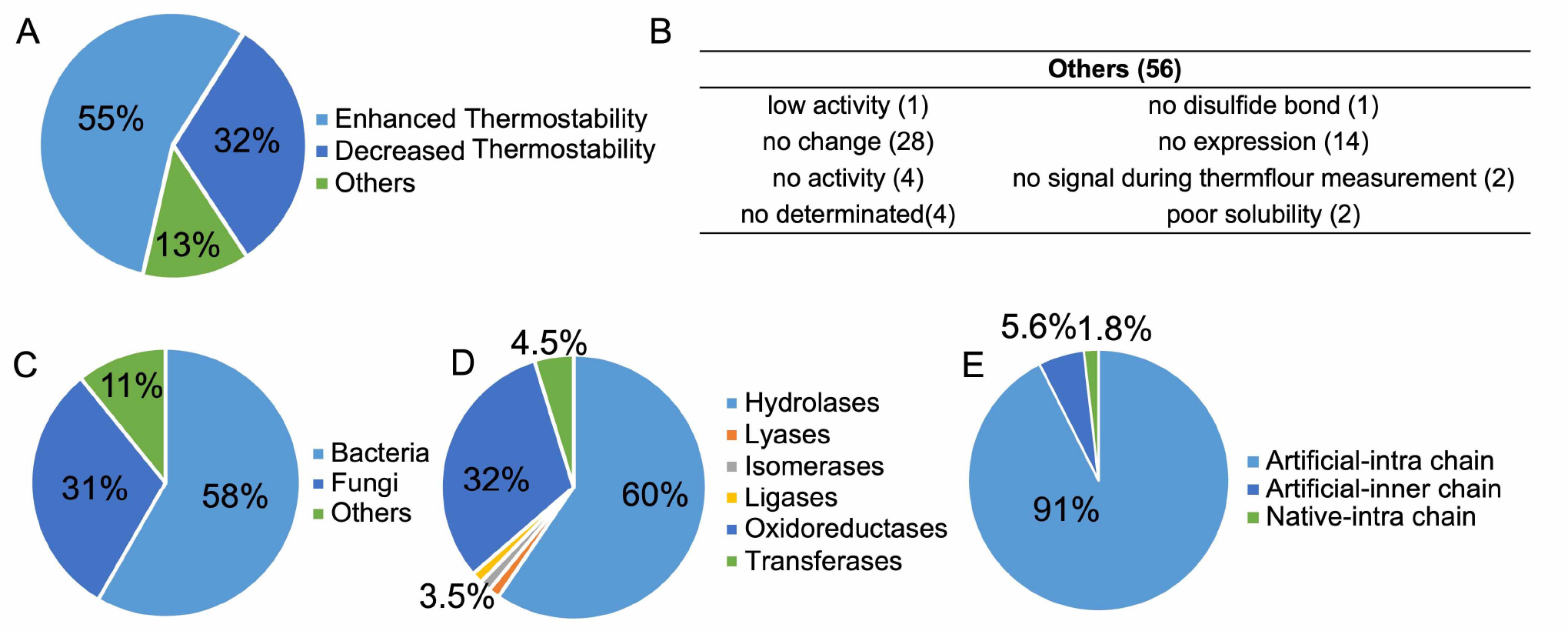
Dataset information. (A–B) Distribution of functional types of SS bonds collected in this work. (C-E) Distributions of source organisms (C), classifications (D) of all enzymes, and type and position of disulfide bonds in enzyme structures within the database, including artificial intra- and inter-chain and native intrachain (E).

Many native SS linkages use alanine and serine mutations for the confirmation of their function, and our data includes eight of these cases. Among the artificial SS linkages, 92.5% are intra-chain (within the same chain) and 5.7% are inter-chain (between different chains) (Fig. 2E). Additionally, 103 enzyme entries were specifically utilized for the analysis of intra-chain SS bonds, while 11 enzymes were specifically employed for the design and investigation of inter-chain SS bonds. In this study, only intra-domain SS linkages were considered in constructing the prediction model.

### Key features derived from the mutation sites

To unearth the critical factors that contribute to the stabilizing or destabilizing effects of engineered SS linkages on enzymes, we extracted features based on the sequence and structure of these engineered enzymes, including evolutionary information, micro-environment information, and other geometric properties of mutation sites. The ranking of the importance scores for the features is shown in Fig. 3, which represents the top 10 essential factors retrieved from the mutation site and the neighborhood context. As can be seen, the most important feature in both models is the mutation-induced Gibbs free energy change. Other features, such as the geometric (distance and dihedral angle), evolutionary (PSSM and BLOSUM62), local interactions (hydrogen bond), residue type, SASA, secondary structure (DSSP), etc., are also very important. Next, a comprehensive analysis was performed to provide a detailed examination of the importance of these features in improving protein thermostability (Fig. 4).

**Figure 3.**
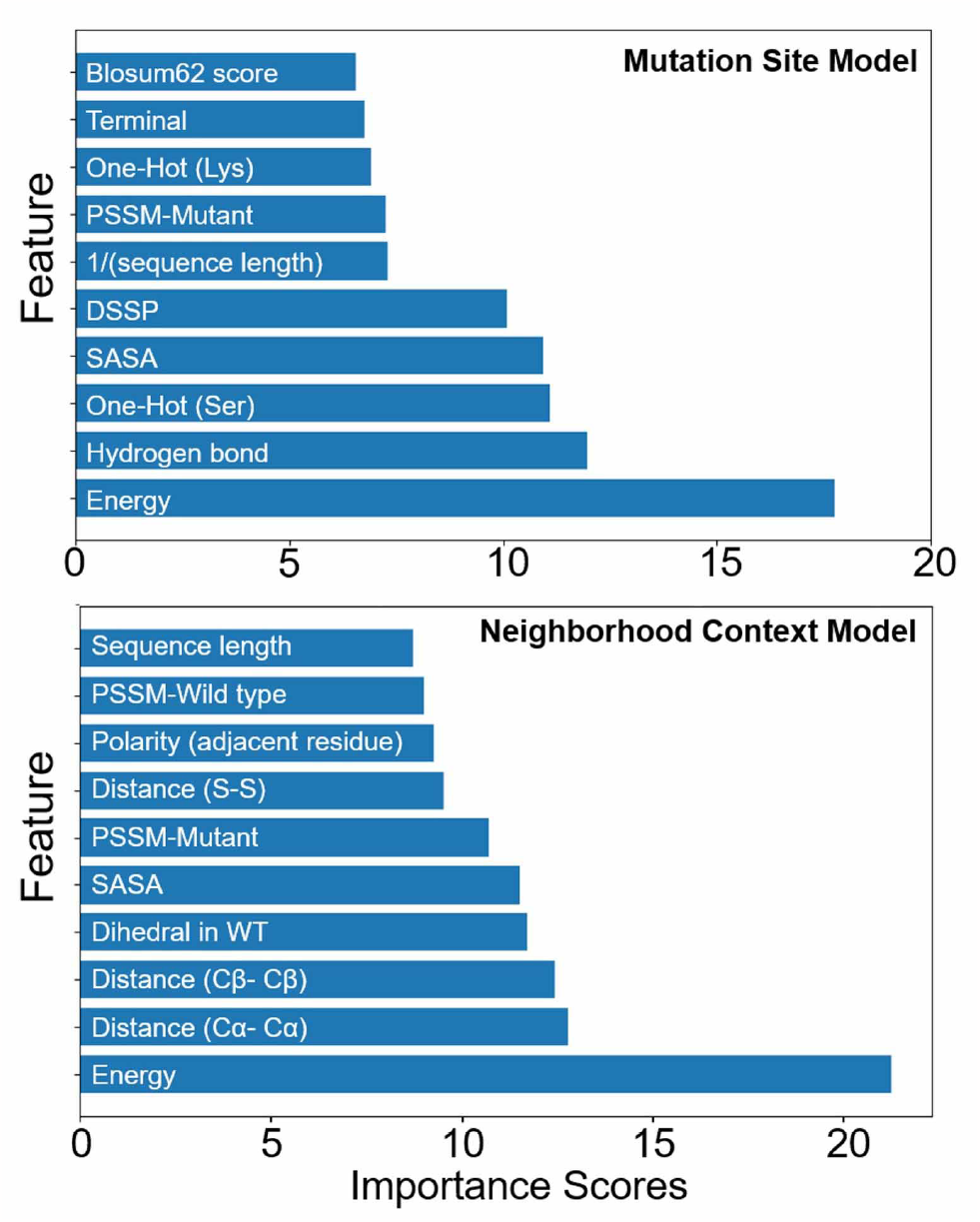
Top ranking features based on their importance. The features at the top represent considering the mutated-site ones only, while those at the bottom consider both mutated-site and neighborhood-context ones.

**Figure 4.**
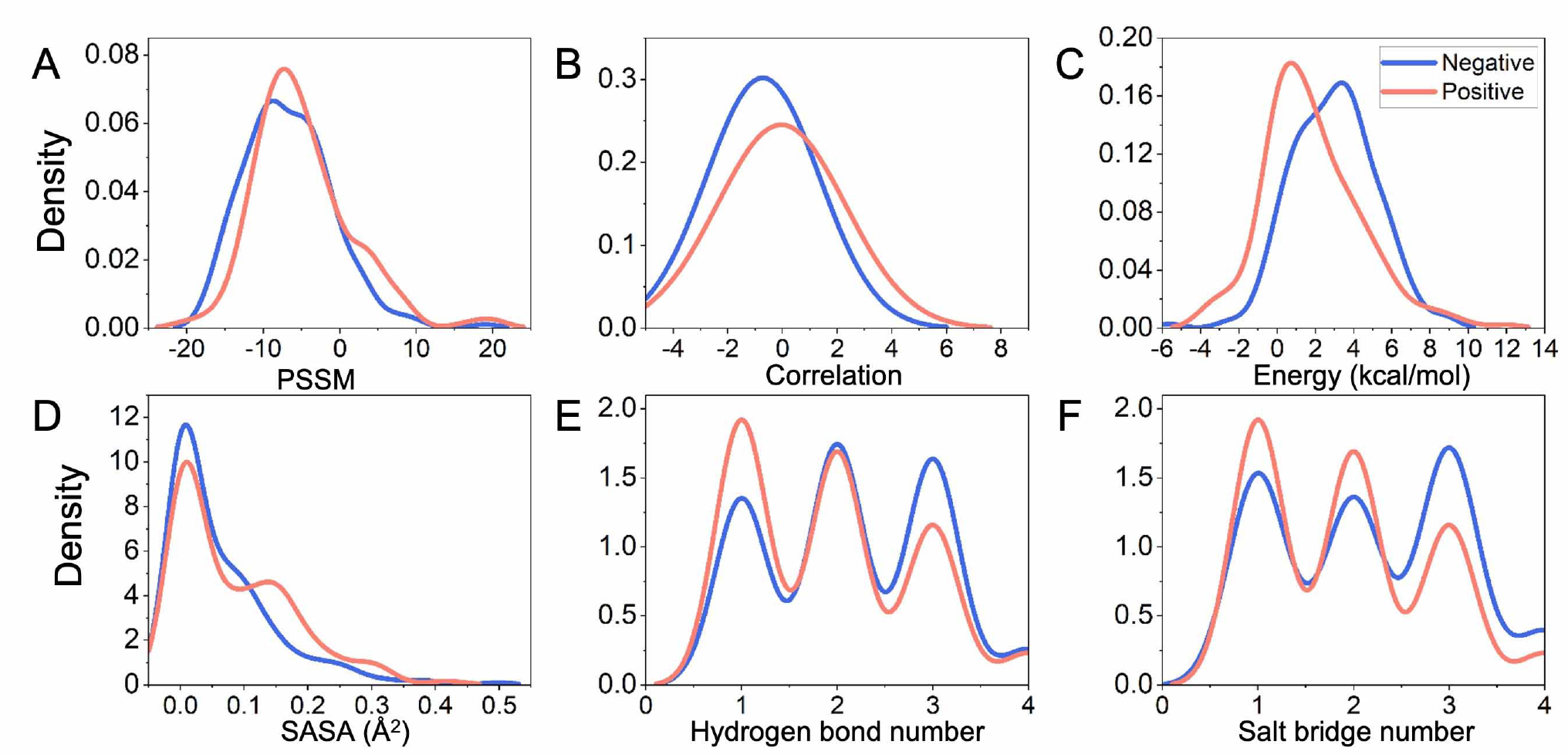
The distribution of key features, including PSSM scores (A), correlation scores between mutation sites calculated using the BLOSUM62 matrix (B), mutation energy upon cysteine mutations (C), the solvent accessible surface area (SASA) of design sites in wild type structures (D), and the non-covalent interactions between mutation sites and their surrounding residues in the wild type structure, including hydrogen bonds (E), and salt bridges (F).

#### Conservation of engineering sites

Zhu et al.[64] found that conservation characteristics played an important role in post-transcriptional translation (PTM), such as disulfide formation. Hence, for the SS-bond design, considering the conservation feature may exert less impact on protein function under double mutations. Herein, we first try to investigate the evolutionary relationship between mutated site pairs involved in SS-bond formation.

Position-Specific Scoring Matrix (PSSM) acts as a widely used computational tool in bioinformatics and protein analysis[43]. Based on multiple sequence alignments (MSAs) of related proteins, PSSM can provide valuable information on the conservation of residues at each position in the alignment. In this study, PSSM was calculated for each sequence and taken into account in multiple mutation prediction algorithms[65]. As shown in Fig. 4A, the PSSM scores were calculated when cystine was present at the engineering sites. For each sample, the score is the sum of the SS-bond sites. In the distribution plot, the PSSM scores of positive samples tend to be larger values. A higher PSSM score indicates a higher likelihood or preference for a particular amino acid at a specific position in a protein sequence.

Furthermore, the correlation scores of mutation site pairs were calculated using the BLOSUM62 matrix[66, 67], a commonly used substitution matrix for the alignment of protein sequences. The BLOSUM62 matrix could be used to evaluate the substitution frequencies between different residues. It is based on extensive protein sequence alignment data and reflects the evolutionary relationships and co-occurrence patterns between different residues. As illustrated in Fig. 4B, the positive SS bonds exhibit a larger correlation of the site pairs. Hence, the positive sites have a slightly stronger evolutionary relationship.

#### The change in Gibbs free energy upon mutations

In protein systems, ΔΔ*G* can reflect the variation in the stability (folding free energy) or binding affinity (between molecules such as antibody-antigen) resulting from a mutation. It represents the energetic contribution associated with altering the protein’s structure, interactions, or conformational dynamics. For rational enzyme design, energy shifts from wild-type to mutant have been taken into account[68, 69]. Many protocols have been developed aiming at calculating free energy differences and have grown in popularity over the past years. Assessment of energy change is typically based on thermodynamic parameters, such as the free energy difference (ΔΔ*G*). A larger positive value indicates that the mutation may lead to enzyme instability, while a smaller value suggests that the mutation may be tolerable.

In this work, the energy for each pair of residues involved in disulfide bonds was calculated using the FoldX suite[70]. As shown in Fig. 4C, distinct variations in peak values can be observed in the distribution curve of changes in energy value for increased thermostability, as well as in other types of data display. The SS bonds with enhanced thermostability are characterized by smaller energy changes, further demonstrating the importance of energy calculations for improving thermostability using SS-bond design.

#### The exposure level of engineering sites

The solvent-accessible surface area (SASA) is a measure that describes the contact extent between residues located at the protein and the surrounding solvent[71, 72]. According to the distribution curves in Fig. 4D, the positive samples show higher values in the distribution of SASA before the introduction of SS bonds, suggesting that incorporating SS bonds at solvent-exposed positions might potentially contribute to enhancing protein stability. However, the introduction of SS bonds may restrict enzyme flexibility and impede conformational changes in the active site, potentially impairing enzyme function. So, caution needs to be taken when designing SS bonds, particularly in proximity to essential functional sites.

#### Non-covalent interactions around mutation sites

To achieve and maintain the stability of the protein structure, a diverse range of non-covalent interactions play a crucial role. These interactions include hydrogen bonds, salt bridges, Van der Waals interactions, and various other types. Here, focusing on the polar interactions, we calculated the native hydrogen bonds and salt bridges around the mutation sites, and compared their distribution between positive and negative samples. As shown in Fig. 4E–F, the strength of native attractive interactions around engineering sites is weaker in the positive data compared to the negative ones. When the introduction of an SS bond fails to improve the thermostability of the enzyme, it suggests a tendency to preserve the original conformation, resulting in a relatively minor impact on its stability. Hence, taking into account native interactions around the wild-type sites, the design process can be strategically optimized to preserve or improve the stability and functionality of the protein, enhancing the probability of achieving the desired modifications with precision and efficacy.

#### Amino-acid type and position of the engineering sites

To determine the impact of residue substitutions on enzyme thermostability, we conducted an analysis of residue frequency distribution specifically at the cysteine mutation sites within each of the two categories Fig. 5A. Alanine has the highest frequency in both categories, and this is consistent with the BLOSUM62 matrix, indicating A has a high substitution probability with C. Interestingly, mutations that are more likely to be associated with improved enzyme thermostability involve conservative replacements, such as S and T, which may have less influence on local polarity and volume[73]. Typically, considerable proteins possess unbounded terminus and the stabilization of these unbounded short regions can greatly improve the overall thermostability of the enzyme. For example, Zhou et al.[74] successfully introduced an N-terminal disulfide to improve the thermostability of a GH11 xylanase derived from *Aspergillus niger*. As illustrated in Fig. 5B, according to our collected data, SS bonds involving terminal residues make a greater contribution to the improvement of thermostability. Hence, when designing disulfide bonds, it is advisable to pay particular attention to the A/S/T residues and terminal regions.

**Figure 5.**
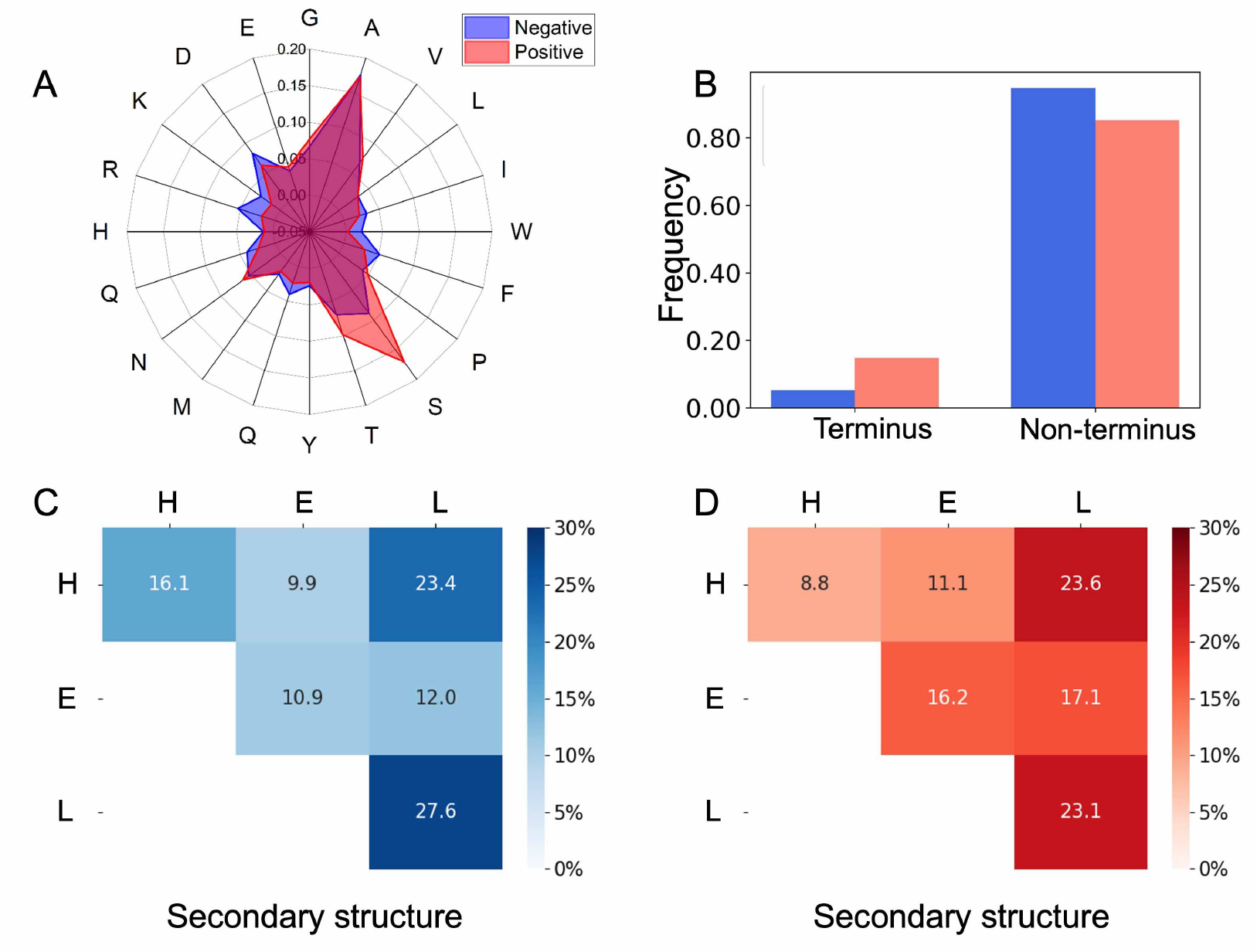
Key characteristics of mutation sites: (A) the residue-type distribution of mutation sites; (B) the location of SS bond design site in terminal or non-terminal, and terminal SS bond is one in which at least one residue is within the first five and last five of the entire sequence; (C-D) heatmap plots of secondary structure information of mutation sites for negative and positive samples, respectively.

#### Secondary-structure preference of the disulfide sites

A common design strategy for thermostability improvement is to rigidify the flexible regions. However, our findings demonstrate that SS bond sites occur preferentially in disordered regions (L), regardless of whether they are positive or negative (Fig. 5C–D). This suggests that the introduction of SS bonds is facilitated by flexible regions. Nonetheless, it is not necessarily true that the rigidification of flexible regions will always lead to improved thermostability. Furthermore, SS bonds, involving stable secondary structures such as the helix and *β*–sheet, exhibit specificity in promoting changes in protein thermostability. For example, the positive samples prefer *β*– sheet, the H-L combination has no discernible difference, while the H-H and L-L ones tend to induce negative results.

### Key features of residues near the mutation sites

The micro-environment formed by different residues plays a central role in the formation of protein 3D structures[75]. As SS bonds contribute significantly to protein structural stability, the micro-environment surrounding an SS bond should also affect its functionality. Most natural SS bonds, preserved throughout long-term evolution, are highly likely to contribute thermostability of enzymes. Analyzing the micro-environment difference between artificial SS bonds collected in the present work and the natural ones might provide clues for improving protein thermostability by introducing extra SS bonds. Our subsequent analysis considers both sequence- and structure-based micro-environments (Fig. 6).

**Figure 6.**
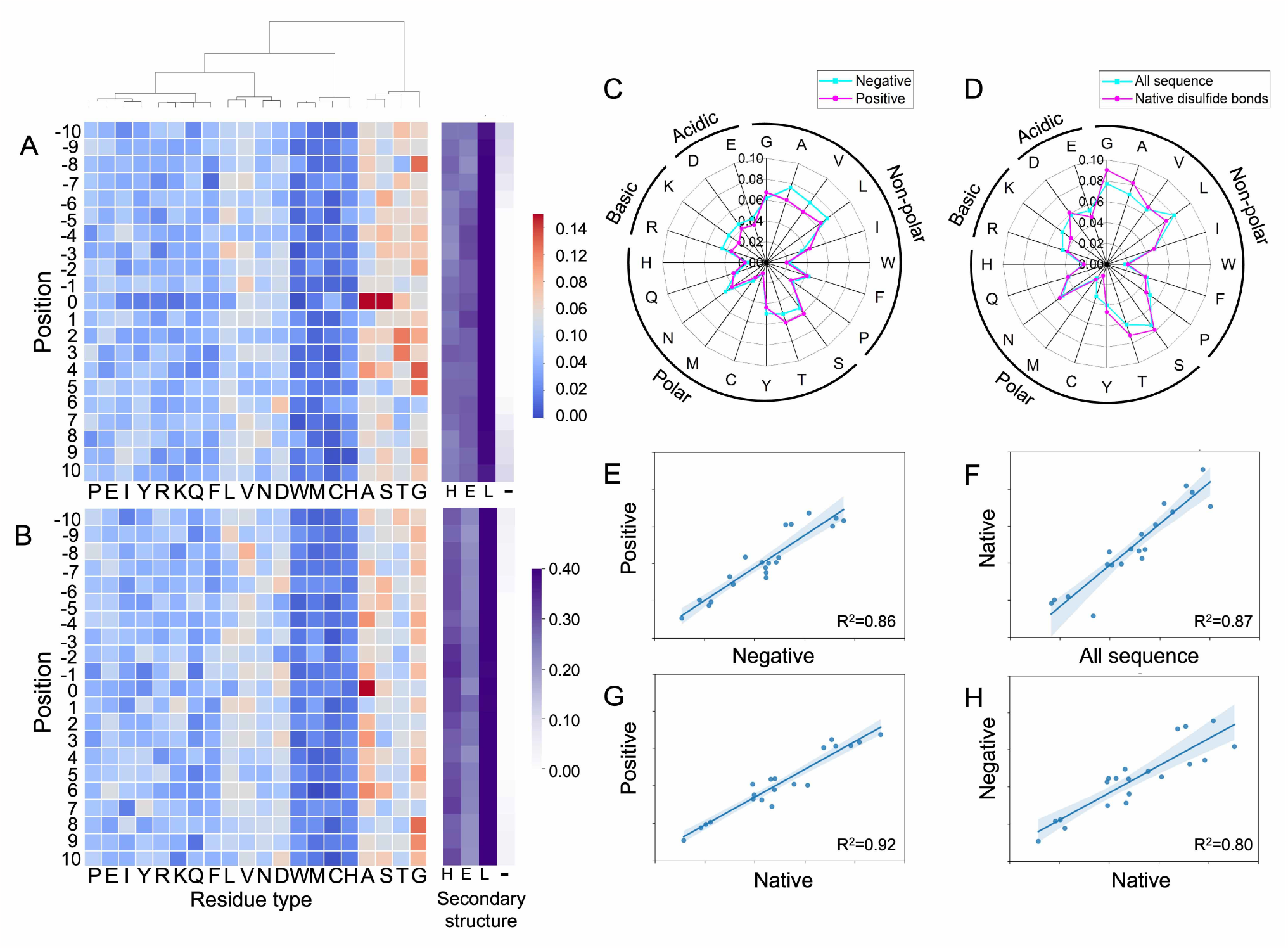
Key features of residues around disulfide-bond designing sites. (A-B) residue-type and secondary structure distribution of sequential neighbor residues for positive (A) and negative (B) data. The residue types of all samples were clustered by probability values. Abbreviations: H, helix; E, sheet; L, loop; –, not predicted. (C) Amino acid preference of spatial adjacency residues around the engineered (negative vs. positive) SS bonds. (D) Amino acid preference of the spatial adjacency residues around the native SS bonds compared to that of all protein sequences. (E-H) Comparisons of two-by-two linear regression curves of four datasets mentioned above. The native disulfide bond information and the corresponding full-length sequences were retrieved from the cullpdb database.

First, we extracted amino-acid sequences of length 21 centered on the mutation sites. Subsequently, a statistical analysis was performed to determine the distribution of residue types at each position. This sequence-based information was compared between positive and negative samples of artificial disulfide bonds. As illustrated in Fig. 6A–B, the distribution of the residue types in both data sets exhibits great similarity and is classified into two primary groups. Residues A, S, T, and G are more prevalent than others, especially A and G. In both datasets, A is the most frequently observed residue type at the mutated sites, while G is popular at other positions, except for the engineering sites. Although S and T are abundant in the positive samples, their contributions are relatively scarce in the negative samples. The frequency of C implies that the formation of SS bonds requires only one mutation along with a natural C. Single mutations generally have less effect than double mutations. Consistent with (Fig. 5C–D), mutation sites located at the beta-sheet are more likely to improve the enzyme thermostability.

Next, the residue distributions in the spatial vicinity of the positive and negative SS bonds were evaluated. As illustrated in Fig. 6C, the distributions of positive and negative data are similar. To verify whether this characteristic is specific to residues in close proximity to disulfide bonds or is related to the overall frequency of amino acids in the protein sequence, we compared the frequency of amino acids in the spatial vicinity of natural disulfide bond sites with the frequency of all amino acids in the protein sequence Fig. 6D. Interestingly, the distribution trends are remarkably similar.

Then a two-by-two linear regression was performed between these results (Fig. 6 E–H). The correlation between the positive and negative datasets is about 0.86, very close to that between natural disulfide bond sites and all sequences (R^2^=0.87). However, the positive data are much more similar to the natural disulfide bonds than the negative ones (0.92 vs. 0.80). Although the local SS-bond sites and overall sequences show a high correlation (R^2^=0.87), there remains a residue preference observed within the specified SS-bond samples. G, with its minimal side chain, predominantly occurs around natural and positive SS bonds. L, with a larger hydrophobic side chain, is less likely to be found near SS bonds. T and S, which are similar to C, frequently appear around positive and natural SS bonds. Based on the aforementioned analysis, utilizing S as a substitution site for C is more likely to yield desired outcomes. Basic amino acids, including K and R, are less frequently observed near positive samples.

### Performance of the binary classification models

To address the current challenge of engineering enzymes, we have developed innovative prediction models focused on the influence of introduced SS bonds on enzyme thermostability. Our model leverages two sets of features: one set based solely on mutation site information and the other set incorporating the information of the mutation site and its surrounding properties. To construct the prediction model, we employed a diverse range of machine learning algorithms, including SVM, XGBoost, Random Forest, Adaboost-Decision Tree (Adaboost-DT), and Adaboost-Random Forest (Adaboost-RF).

Initially, we extracted a comprehensive set of features based on the mutation sites. These features encompass crucial information such as mutation type, conservation information, structure information, and so on. To ensure optimal utilization of these features, we employed advanced feature engineering techniques including feature selection and normalization. In addition, we considered the properties surrounding the mutation sites. We extracted contextual features that encompassed sequence information of neighboring bases, structural characteristics, and other relevant attributes. These contextual features provided a more comprehensive understanding of the mutations. Based on the two sets of features, our prediction model was constructed using various machine learning algorithms.

To evaluate the performance of the models, we assessed key evaluation metrics such as accuracy, recall, F1 score, specificity, AUC, and precision. The results (Table 1) support the efficacy of our proposed model in predicting the causal relationship between an introduced SS bond and the change in the thermostability of the protein. The highest accuracy is 0.714, shown in the mutation site model using Random Forest and Adaboost-DT, and in the neighborhood context model using Adaboost-DT.

**Table 1.**
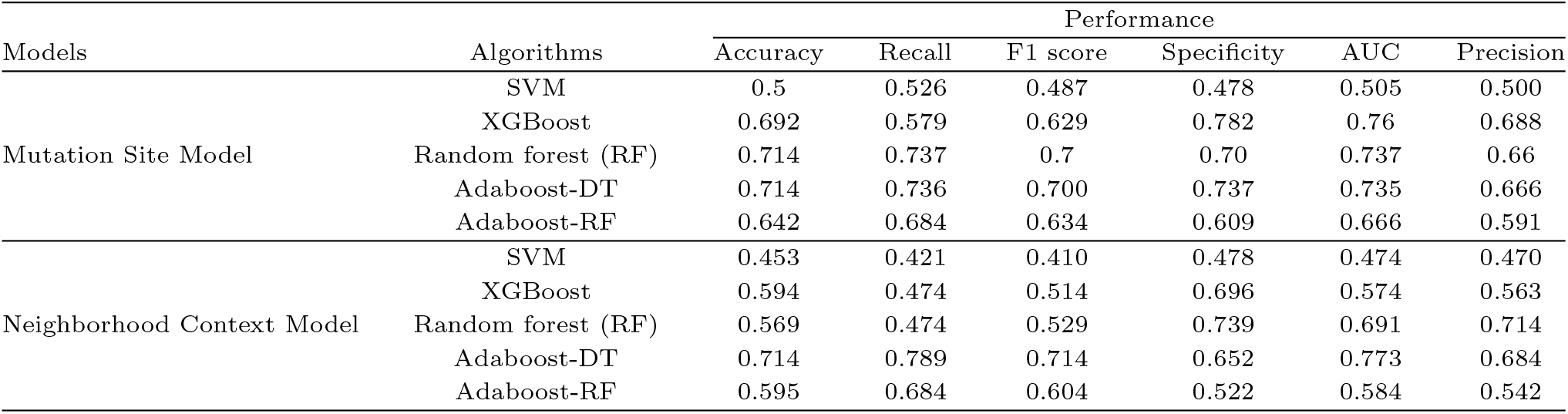
Prediction performance by different methods.

In particular, among the Mutation Site Models, RF exhibited superior performance on multiple metrics, and the AUC score is 0.737, while the Neighborhood Context Model excelled with an AUC of 0.773 using Adaboost-DT. These findings suggest that information-based enrichment can be beneficial for the ability enhancement of the model to discriminate between positive and negative samples.

Above all, our prediction models integrate the characteristics of the mutation sites and the surrounding properties, providing satisfactory results through the utilization of various machine-learning algorithms. The models constructed in this study may serve as a valuable reference for further advances in sequence- and structure-based prediction tasks.

## DISCUSSION

### The superiority of AF2 on SS-bond prediction

Recently, deep learning algorithms such as AF2 predict three-dimensional protein folds with great certainty[69, 79]. The research conducted by Patrick Willems et al.[79] demonstrates a high level of consistency 96% between the SS bonds predicted by AF2 and those observed in X-ray and NMR protein structures. Traditional disulfide prediction algorithms perform predictions mainly based on geometric distances in 3D structures, which could potentially lead to the loss of valuable data. Given the incompleteness of the structure database[22, 80] and the high precision of AF2, we submitted sequences collected from the literature to fastAF2[34, 35] for the prediction of the structure of wild-type enzymes and mutants. For our dataset, the distances of the C*β* atoms in the WT and the sulfur atoms (S-S) in the mutants were calculated between the designed disulfide sites (Fig. 7A-B). Results suggest that some C*β* -C*β* distances are far beyond reasonable limits for the formation of SS bonds in WT structures, while some pairs exhibit a remarkable decrease in the S-S distance, especially those with enhanced thermostability. For some items, the designed sites in wild-type structures do not meet the geometric demands of SS bridges, whereas SS bonds were successfully introduced into these structures. Surprisingly, these long-distance engineering sites are located at the protein terminus (Fig. 7C–F), allowing the relatively unfettered terminus to immobilize on the protein surface, thus inducing conformational rigidity upon the introduction of SS bonds. To some extent, with the utilization of AF2, the accuracy of SS bond prediction can be enhanced, leading to an improvement in the enzyme thermostability. Therefore, during disulfide design, terminal engineering sites may serve as preferred candidates, as they are more likely to increase enzyme stability. Meanwhile, AF2 shows the potential to predict SS bonds formed by residues that are spatially distant in enzymes. In summary, terminal SS bonds can be designed and subsequently filtered using our model, presenting a highly efficient strategy for improving enzyme thermostability, especially for those lacking experimentally determined structures. Additionally, the introduction of such non-native cross-links into protein structures may unbalance the complicated structure-function relationships, and the mechanisms need in-depth investigation for accurate digging.

**Figure 7.**
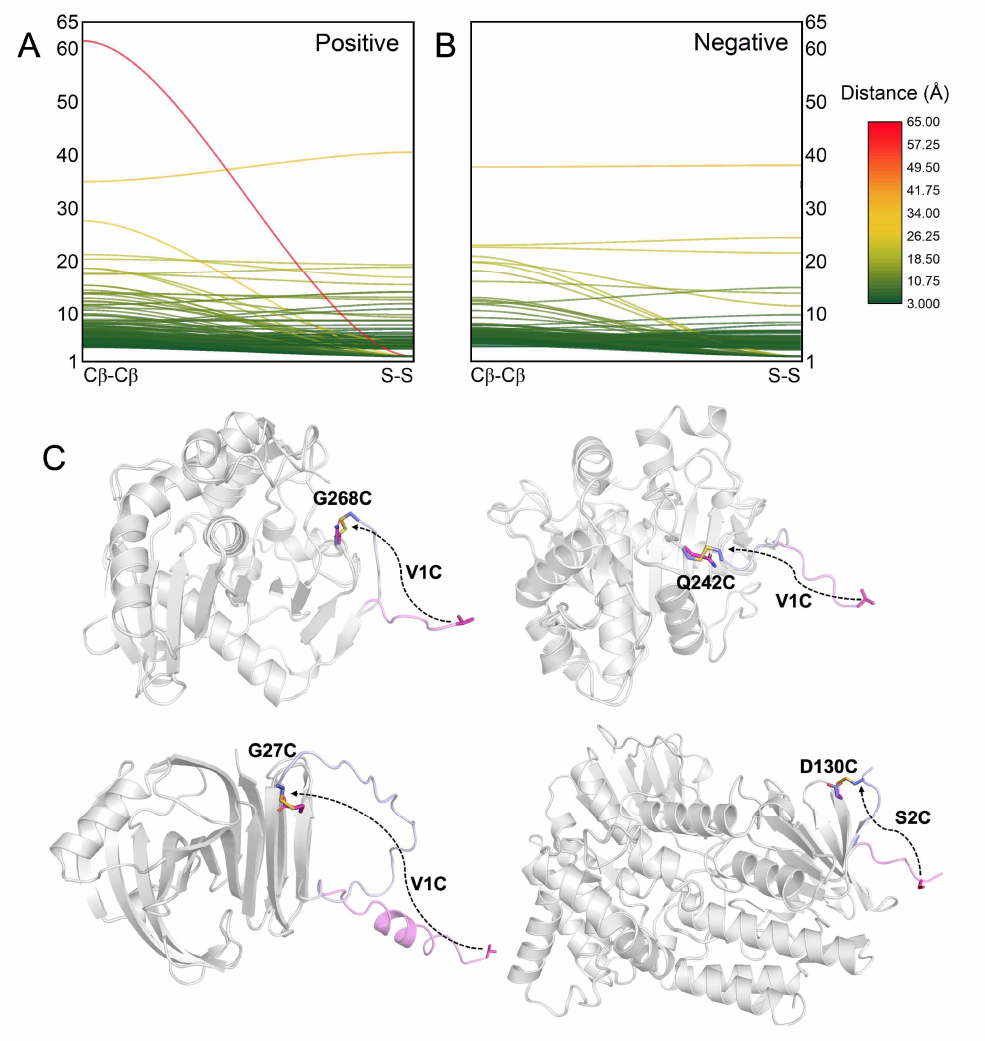
The side-chain distances between the engineering sites before (C*β* -C*β*) and after (S-S) the introduction of SS bonds (A) and representative examples of successfully engineering SS bonds captured by AF2 (B). (C-F) Structural superposition between AF2-predicted structures for wild-type and mutant forms[77, 78, 74]. The mutants containing new SS bonds are assigned numbers in the database, indicated at the bottom right corner of each figure.

### The importance of co-evolution information

Rational and effective disulfides should be considered at the interrelated residue pairs during evolution, which may reduce the adverse effects on protein folding or conformation. In the aforementioned analysis, we found a certain positive correlation between the conservation of residues involved in the formation of SS bonds and the enhancement of protein thermostability. Furthermore, the co-evolutionary patterns of the residue pairs should also be favorable for the engineering of disulfide bonds. To verify this hypothesis, we used the thermostability modification data derived from Wang et al.[81] as examples to test the potential relationship between the co-evolution and thermostability of disulfide residue pairs. A Python framework named EVcoulpings proposed by Thomas A Hopf et al.[82] was employed. This tool can perform co-evolutionary sequence analysis for de novo prediction of the structure and function of RNA, proteins, and protein complexes. Evolutionary couplings (EC) between sites can be identified with the assistance of the EVcouplings software, which also corresponds to physical contacts in the 3D structure of the molecule. As shown in Table 2, there is a positive correlation between EC scores and changes in thermostability for the disulfide mutant. Although the observed phenomenon in our dataset may not always follow that pattern, for the design of the SS linkage, we still need to consider the conservation and co-evolutionary information of the residue pairs as key factors.

**Table 2.** Ev-coupling score and thermostability function of SS bond from Endo-polygalacturonase Talaromyces leycettanus JCM 12802[76].

| Source | Mutation pairs | Thermostability | Ev-coupling score |
| --- | --- | --- | --- |
| Endo-polygalacturonase from *Talaromyces leycettanus* | T316C/G344C | Remarkably Enhanced | 16.991 |
|  | N253C/G282C | Little Enhanced | 12.522 |
|  | G220C/S249C |  | 12.406 |
|  | I99C/S140C |  | 3.415 |
|  | N54C/N78C | Decreased | / |
|  | E86C/T127C |  | / |
|  | K128C/V152C |  | / |
|  | T182C/N202C |  | / |
|  | S211C/D238C |  | / |

### The current stage and challenge of engineering SS bonds for enzyme thermostability improvement

SS-bond design in response to thermostability modification of enzymes is of great importance. Many models have been proposed for the prediction of disulfides, such as DBD2[83], SSBOND-Predict[84], and Yosshi[25]. DBD2 is the second-generation design tool derived from the SS bridge design (DBD)[83] that incorporates conformational constraints and introduces the B-factor for the design of SS bonds, while SSBOND-Predict is a neural network-based model to predict amino acid pairs to construct engineered SS bonds[84]. Another application, Yosshi, integrates evolution information through multiple sequence alignment (MSA) for SS-bond prediction[25]. All of these existing computational tools have made significant progress in the identification of potential disulfide-bond sites. However, at the practical view, achieving the objective of enhancing thermostability remains a challenge because of inadequate consideration and additional efforts are required to improve the accuracy and reliability of the prediction to bridge the existing gap.

Now, we attempt to represent an extended design strategy that begins with the existing SS-bond prediction tools and subsequently incorporates the use of our models to refine the selection of potential variants, with the aim of improving enzyme thermostability (Fig. 8A). To achieve this, a set of potential SS-bond sites can be generated in the target enzymes using existing prediction tools. Given the portability of DBD2 and SSBONDPredict, we chose them to predict the SS-bond sites. Although the performance of DBD2 in our test data set is slightly better than that of SSBONDPredict, it is important to note that the accuracy of both tools remains relatively low (Fig. 8B). Some disulfide bonds might not be hooked due to the structural rigidity of proteins. Regarding the SS bond detected by DBD2, less than half of them can positively affect the thermostability of enzymes, indicating the need for further screening. The potential sites are then subjected to enzyme thermostability prediction to screen the SS-bond candidates using our model, which is specifically designed to detect intricate patterns and characteristics related to SS bonds and enzyme thermostability.

**Figure 8.**
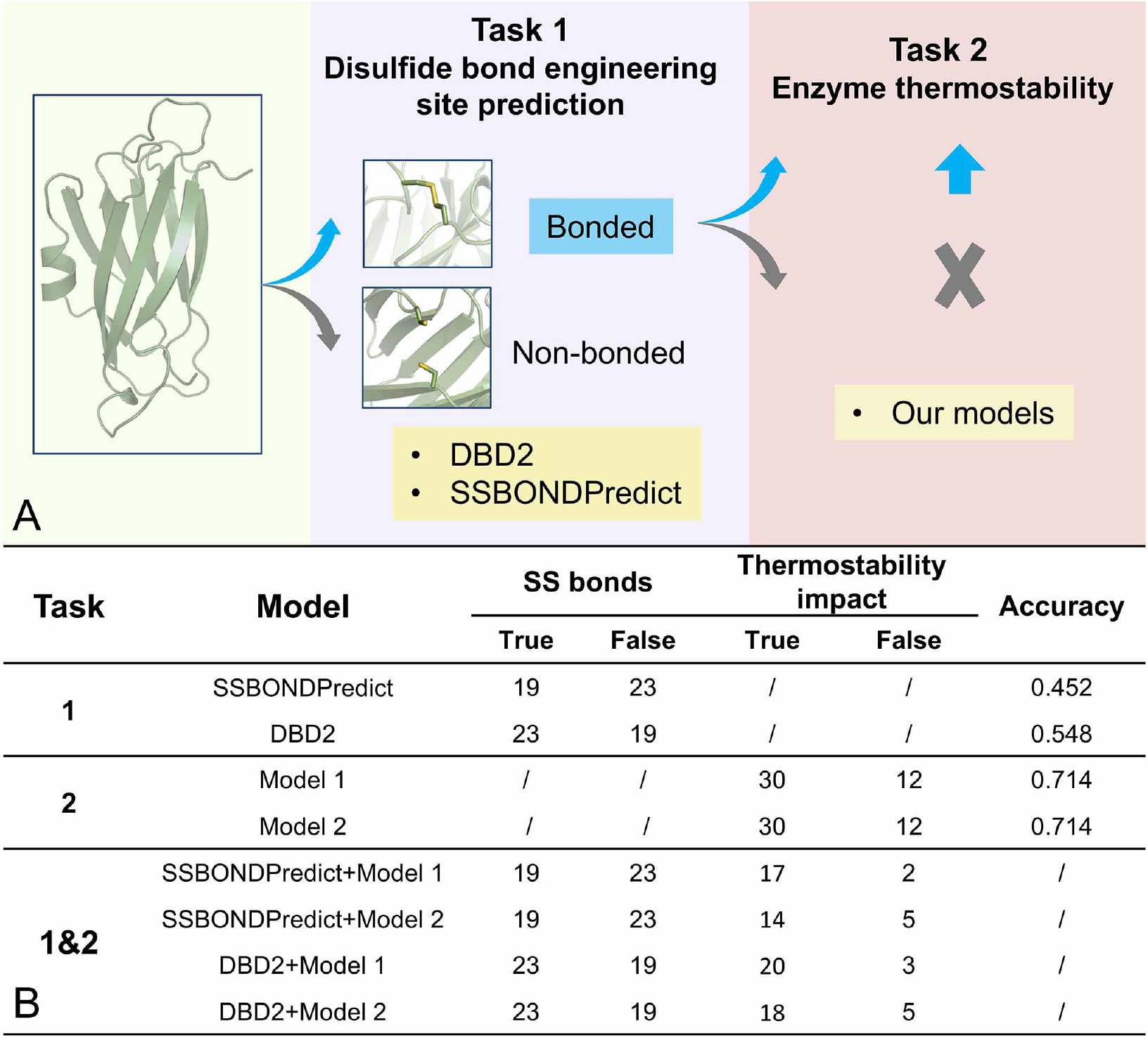
An extended design strategy for SS bond design incorporating the existing tools and our models. (A) Relationship between our work and other existing SS bond prediction tools. Potential SS bonds can be predicted using existing tools, such as DBD2 and SSBONDPredict, while the aim of our models is to evaluate whether the introduction of an SS bond can improve the thermostability of the enzyme. The integrated application of these two kinds of tools offers the potential for enhancing enzyme thermostability by facilitating the introduction of SS bonds. (B) The performance of the existing models and our models on the testing dataset, including individual and combined strategies.

Furthermore, by analyzing the test set in this study, our extended strategy produces a set of predictions that present high positive results, demonstrating high precision in identifying positive SS-bond sites, greatly improving the target of the original predictions. This accuracy is significant for enzyme engineering, as it provides a more reliable method to guide enzyme design and optimization, further advancing the development of biotechnology and industrial applications.

## CONCLUSION

SS bridges, as a feasible and powerful modification strategy in enzyme engineering, are often introduced to enhance the thermostability of enzymes[85, 86, 87]. But sometimes the results are not always as expected[29, 88]. Here, a manually curated database of enzymes with experimentally determined thermostability data upon the introduction of SS bonds, including inter- and intra-chain SS bridges, thus was prepared. Machine learning models were constructed based on the intra-chain data to introduce SS bonds more accurately and efficiently. Sequence- and structure-based features, as well as conservative and energy ones, were extracted. The conservative and energy features both have evidently different distributions between the two categories. Overall, this suggests that the prediction model presented here can be effective in identifying possible residue pairs to introduce SS bonds to improve enzyme stability with fewer failed attempts. Also, compared to existing tools, our model performs better in SS bond enhancement. In summary, our work, featuring an improved model for more accurate and efficient prediction of SS bonds, will significantly facilitate enzyme engineering for industrial applications.

## SUPPLEMENTARY DATA

All data are incorporated into the ThermoLink online server and can be accessible at guolab.mpu.edu.mo/thermoLink. Other data is available upon reasonable request.

## Author contributions statement

J.G. and L.Z. conceived and designed the study. R.X., J.G., and L.Z. collected and analyzed the data. J.G., R.X., and Y.Y. designed and built the online server. J.G., L.Z., R.X., Y.Y., Y.W, M.X, Z.W., S.W., and W.L. wrote and revised the manuscript.

